# Rewriting protein alphabets with language models

**DOI:** 10.1101/2025.11.27.690975

**Authors:** Lorenzo Pantolini, Gabriel Studer, Laura Engist, Ieva Pudžiuvelytė, Florian Pommerening, Andrew Mark Waterhouse, Stefan Bienert, Gerardo Tauriello, Martin Steinegger, Torsten Schwede, Janani Durairaj

## Abstract

Detecting remote homology with speed and sensitivity is crucial for tasks like function annotation and structure prediction. We introduce a novel approach using contrastive learning to convert protein language model embeddings into a new 20-letter alphabet, TEA, enabling highly efficient large-scale protein homology searches. Searching with our alphabet performs on par with and complements structure-based methods without requiring any structural information, and with the speed of sequence search. Ultimately, we bring the exciting advances in protein language model representation learning to the plethora of sequence bioinformatics algorithms developed over the past half-century, offering a powerful new tool for biological discovery.

## 1 Introduction

Protein sequence alignment has been a cornerstone of bioinformatics for decades, with its use growing significantly alongside expanding databases and deep learning advances in structure and function prediction [1–4]. Methods like BLAST [5] (Basic Local Alignment Search Tool) revolutionised database searching by using heuristics to find regions of local similarity, offering a fast, albeit less sensitive, alternative to exhaustive dynamic programming algorithms. Tools like MMseqs2 [1] (Many-against-Many sequence searching) led to further massive speed improvements by introducing highly efficient indexing techniques and *k*-mer matching strategies to accelerate the initial search phase, making it possible to compare sequences against massive protein databases in minutes. The increasing availability of GPUs has also played a role in optimisations, with MMseqs2-GPU [2] now achieving high speeds without the need for *k*-mer matching. However, sensitivity remains a critical objective in sequence comparison, serving an essential role in both functional annotation and structural modeling when close homologs are scarce.

Simultaneously, the recent years of access to unprecedented amounts of high quality predicted structures by methods such as AlphaFold2 [3] has accelerated innovation. The fact that protein structure is more conserved than sequence has driven efforts to incorporate structural information into sequence comparison frameworks. For example, Foldseek uses a structure-based alphabet called 3Di (Three-Dimensional Interaction) in a fast and sensitive sequence alignment approach to detect remote homology between sequences with similar folds [6]. More recent works such as ESM3 [7] and FoldToken [8] have also used structural tokenization, but these typically create thousands to hundreds of thousands of tokens with the aim of faithful structure reconstruction, too sparse to use in sequence comparison frameworks. However, most sequence databases lack corresponding high-confidence structural models. For instance, billions of protein sequences are available in databases like UniProt [9], MGnify [10] and now LOGAN [11]. In contrast, far fewer experimentally determined protein structures exist in the PDB [12]. While predicted structure databases like the AlphaFold Database (AFDB) [13] and the ESM Atlas [14] help close this gap, a large fraction of their residues are structurally repetitive or have low confidence. In addition, storing and searching databases of millions of predicted structures requires significant resources and compute. This disparity limits the applicability of structure-based methods across the full breadth of sequence space. Furthermore, structural comparison is suboptimal for rapidly evolving proteins with poor predictions and for proteins with extensive disordered regions that complicate structural alignment.

In parallel, protein language models (pLMs) have emerged [14–16], excelling at remote homology detection by generating high-dimensional embeddings that capture complex evolutionary and structural information. The power of these models lies in representation learning: their ability to compress the vast sequence space into dense, numerical vectors (embeddings) where evolutionary and functional relationships are represented. Recently, pLM embeddings have been directly harnessed for remote homology detection. This includes using sequence-level aggregated representations for fast comparison, such as in ProtTucker [17] and TM-vec [18], where the distance between two sequence embeddings is used as a proxy for evolutionary distance. Other methods, such as EBA [19] (Embedding-Based Alignment) and pLM-BLAST [20] leverage the per-residue embeddings to generate sequence alignments. While aligning these continuous representations at the residue-level is highly sensitive, it is no match to the speed and scalability of discrete sequence alignment.

Here, we trained a shallow neural network to discretise pLM embeddings, generating a new 20-letter alphabet, TEA (The Embedded Alphabet), to express protein sequences. Our new alphabet retains the exceptional remote homology detection of pLMs while enabling large-scale comparisons using highly optimised alignment tools like MMseqs2. It performs on par with structure-based approaches and complements them in cases where structural data is suboptimal. Furthermore, TEA provides a confidence metric, offering a valuable estimate of prediction reliability for downstream analysis.

## 2 Results

### 2.1 Rewriting the language of life with contrastive learning

Embedding discretisation can be achieved in various ways. A straightforward approach involves predicting a probability vector, sized to the desired alphabet, and then converting it to the character with maximum probability. This method naturally aligns with the typical training objective of protein language models (pLMs), which often reconstruct protein sequences from embeddings by predicting token probability distributions (including amino acids). However, by modifying the training objective, we can learn a new representation that forces alignment of residues onto the same characters, thereby enabling the comparison of remote homologs.

To achieve this, we trained a shallow discretisation head (depicted in Figure 1A) to convert pLM embeddings into logit representations, and subsequently character probabilities, which are then transformed into discrete characters. This head was trained using a contrastive learning objective designed to force similar character probabilities for structurally aligning residues from homologs, while pushing dissimilar probability vectors for non-aligning residues. As described in Figure 1A, we created residue triplets from structurally aligning residues: an anchor, a positive (the anchor’s structurally aligning counterpart), and a negative (a randomly selected residue from a 5-residue window around the aligning position). We also incorporated losses favouring a uniform character distribution, and low Shannon entropy across the predicted probabilities. The training paradigm is described fully in Section 6.2. This training forces the network to produce characters that result in higher sequence identity for residues sharing similar structural characteristics, producing an alphabet that enables comparison of remote homologs. As shown in Figure 1B, while the pairwise amino acid sequence identity of SCOPe40 alignments is low, the TEA sequence identity is much higher. This holds true even for entire protein folds not included in the training set, demonstrating the generalisability of our alphabet to sequences and folds not seen by our model.

**Fig. 1.**
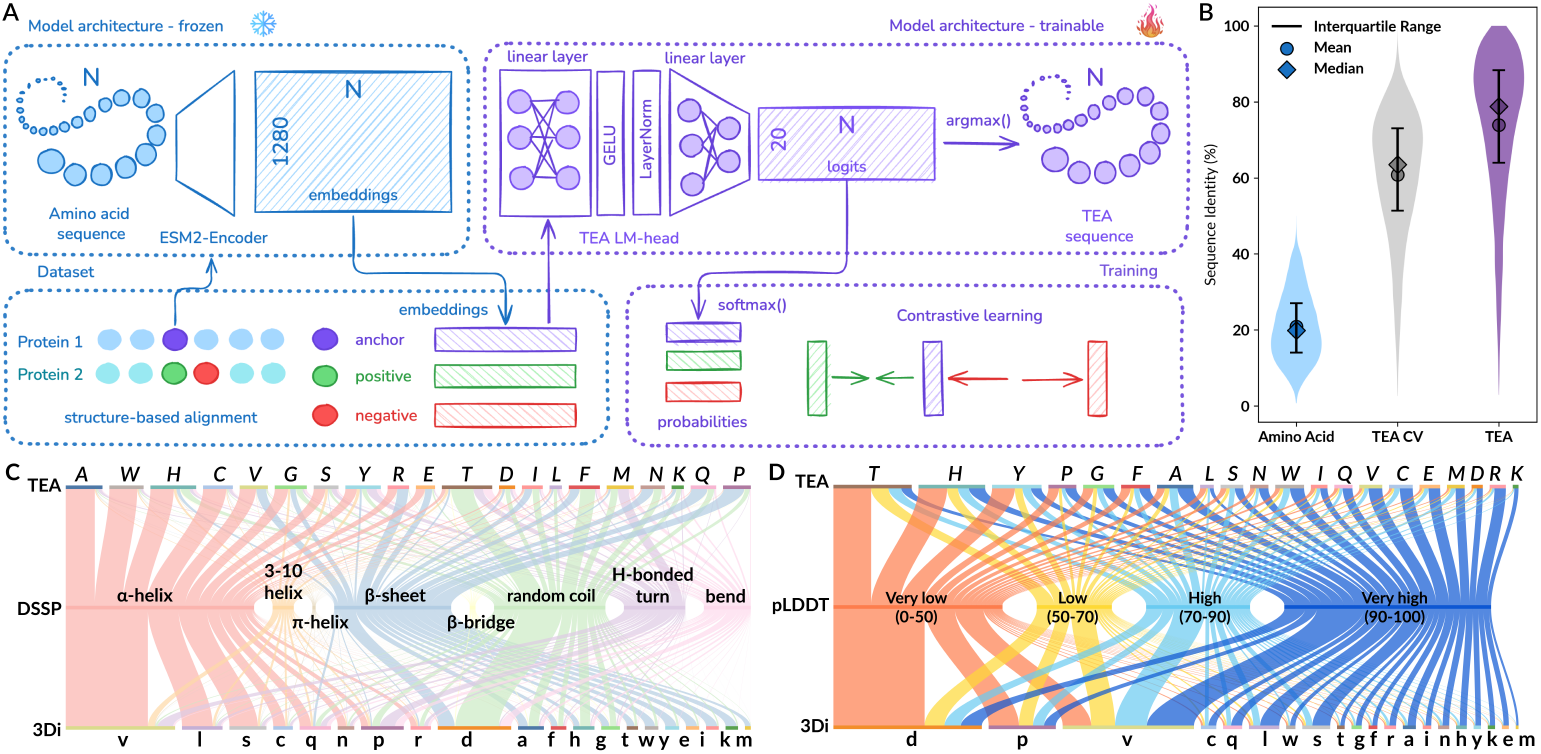
Model and training. **A)** Overview of the model architecture and training regimen. **B)** Sequence identities derived from 24,814 structural alignments (max. 100 per superfamily) of proteins in the same SCOPe family having sequence identity *<*40%, for amino acids and TEA converted sequences. TEA CV represents a 4-fold cross-validation across SCOPe40 folds described in Section 6.1.1, where sequence identities are reported for the alignments from the held-out fold. **C)** A Sankey diagram depicting the distributions and comparisons of (1) TEA characters, (2) DSSP secondary structure labels, and 3) 3Di characters for 15,128 proteins from SCOPe40 and 10,000 proteins from AFDB with average pLDDT*>*90. **D)** A Sankey diagram depicting the distributions and comparisons of (1) TEA characters, (2) AFDB pLDDT, and 3) 3Di characters for 40,000 proteins from AFDB with 10,000 from each pLDDT bin.

#### 2.1.1 TEA as an alternative to 3Di

Given the similarity in approach of discretising continuous data into a 20-letter alphabet, we sought to ascertain whether we learn similar patterns as the 20 characters representing the 3Di alphabet underlying Foldseek, built by vector quantization of protein structural information [6]. TEA characters correlate strongly with secondary structure annotations (Figure 1C rows 1 and 2)^1^. 3Di, like TEA, also has heavy association with secondary structural elements (Figure 1C rows 2 and 3) but differs in terms of distribution and numbers of characters assigned to the different elements. In fact, a majority of residues in helices are assigned to the 3Di-v and loops to 3Di-d, which are known to be overrepresented in larger datasets [21], while multiple letters in TEA all represent helices (*A, W, H, C, G* etc.), sheets (*P, Q, K*, etc.) and loops (*T, F, M* etc.). As shown in Figure 1D (rows 2 and 3), 3Di by definition relies on the quality of the underlying structural data used for construction with most low pLDDT residues manifesting as disordered loops and thus mapping to 3Di-d (54%). Meanwhile, TEA characters are more uniformly spread Figure 1D (rows 1 and 2), with only 29% of low pLDDT residues mapping to *T, F*, and *M*. Some examples of common fully conserved TEA motifs of length 17 are illustrated in Supplementary Figure S1. These motifs are well-conserved in secondary structural elements despite low conservation at the amino acid level, and the motifs themselves range from being found across different protein folds to only being found within a certain protein family. This highlights a promising opportunity for motif discovery to detect structural or functional similarities.

### 2.2 TEA searches reach structural accuracy at a fraction of the cost

We benchmarked our alphabet’s remote homology and structural similarity detection performance. We first used MMseqs2 with sequences converted to our TEA alphabet and a custom substitution matrix (see Section 6.3). As shown in Supplementary Figure S3, and in contrast to the BLOSUM matrix, many off-diagonal elements of the TEA substitution matrix are highly negative, indicating orthogonal characters. As each of the three alphabets, amino acid (AA), 3Di, and TEA come with their own associated substitution matrix, the substitution scores can be combined during the alignment stage to potentially boost performance (see Section 6.5 for implementation details).

As Figure 2A-B shows, TEA (purple circles) achieves sensitivity comparable to Foldseek [6] (orange crosses) and raw embedding-based alignment [19] (green triangles) for detecting relationships at both family and superfamily level on the SCOPe40 benchmark [6]. MMseqs2 and Foldseek employ a sophisticated similar *k*-mer generation technique by default, enabling sensitive searches at the expense of speed. Using exact *k*-mer matching insTEAd causes a drastic decrease in performance when running MMseqs2 with the 3Di alphabet and substitution matrix (yellow diamonds, solid vs. dashed lines). Conversely, TEA yields identical performance in both exact and sensitive modes (purple circles), and reaches structure-level sensitivity with exact *k*-mer matching, underscoring the intrinsic representation power of the alphabet. Supplementary Figure S2 shows the performance of an exhaustive all vs. all alignment without any pre-filtering step for different alphabets and combinations of alphabets. The performances of 3Di (yellow diamonds), 3Di+AA (light yellow pentagons) and Foldseek (orange crosses) clearly demonstrate that amino acid integration and further ranking optimisations are all crucial for Foldseek performance. Foldseek ranking optimisations include, for example, compositional bias correction and reverse bit score addition to suppress FP alignments, and multiplying the bit score with alignment LDDT and TM-score to obtain a structural bit score for ranking [6]. Furthermore, the results from a 4-fold SCOPe fold-level cross-validation (in gray) confirm that TEA remains competitive with structure-based searches even on unseen protein folds. Excitingly, the combination of 3Di with TEA (light purple diamonds in Figure 2A) performs better than either individually, pointing to complementarity between the two representations. We leave the development of search tools making use of such combinations for future work. In all search results reported from this point, unless otherwise specified, TEA is run with exact *k*-mer matching mode.

**Fig. 2.**
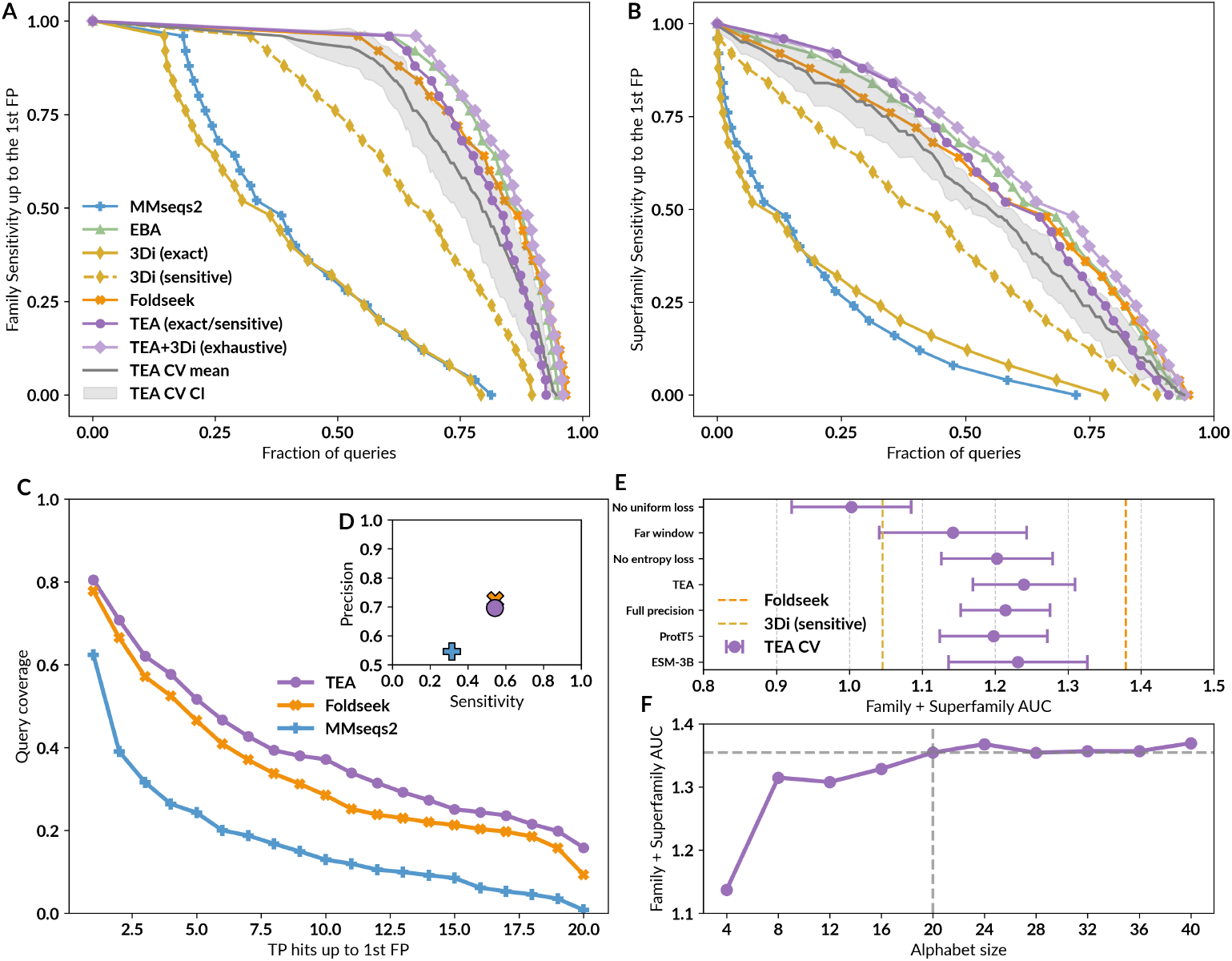
TEA reaches similar sensitivities as structural aligners. **A)** Cumulative distributions of sensitivity for homology detection on the SCOPe40 database of single-domain structures. TPs are matches within the same family; FPs are matches between different folds. Sensitivity is the area under the ROC curve up to the first FP. **B)** Same as A with TPs as matches within the same superfamily. **C)** Search sensitivity of one hundred queries against 56,574 multi-domain, full-length AlphaFold2 protein models. Per-residue query coverage (y axis) is the fraction of residues covered by at least x (x axis) TP matches (LDDT > 0.6) ranked before the first FP match (LDDT < 0.25). **D)** Alignment quality for alignments from E, averaged over the top five matches of each of the 100 queries. Sensitivity = TP residues in alignment/query length; precision = TP residues/alignment length. **E)** Sum of cross-validation AUCs of family and superfamily curves (mean and confidence intervals) for various model ablations (see Section 6.2.1), with the AUC for 3Di (sensitive) and Foldseek shown as yellow and orange dotted lines respectively. CV results in A-C are shown only for held-out SCOPe folds. **F)** Sum of family and superfamily curve AUCs shown for models trained with different alphabet sizes. The final 20-character TEA model is indicated with dotted lines.

Moving beyond single domain proteins, Figure 2C-D presents search sensitivity and alignment quality results for full-length multi-domain AFDB proteins. To avoid predicted disorder, we restricted the dataset to proteins with an average pLDDT > 90. All assessments were reference-free, based solely on the LDDT of the resulting alignments. Across both benchmarks, TEA demonstrates accuracy similar to Foldseek, and much higher than MMseqs2 using the standard amino acid alphabet.

#### 2.2.1 Ablations

We justify our model and training choices through ablations run on the 4-fold SCOPe fold CV, with results depicted in Figure 2E. Enforcing uniform character usage through an explicit uniform loss (Equation 3) improves performance, as otherwise training favours fewer characters being utilised and thus reduces representation power (*No uniform loss* in Figure 2E). Crucially, moving the window further for selecting non-aligning (negative) residues for training worsens performance (*Far window*), as the embeddings of the obtained negatives become much easier to separate from the anchor and positive residues. Neighbouring residues, on the other hand, have very similar residue contexts and thus very similar embeddings to the anchor residue, forcing the model to learn a more nuanced separation between them, leading to better generalisation. Removing the explicit entropy loss term (Equation 2) also slightly worsens performance (*No entropy loss*). Using embeddings from a 4-bit quantized [22] ESM2 model did not show much difference compared to using full precision embeddings (*Full Precision*), thus unlocking fast and memory-efficient TEA sequence generation on consumer-grade GPUs even for longer proteins (0.05-0.5 seconds/sequence depending on length). Using embeddings from the 3 billion parameter ESM2 model (*ESM-3B*) or using the ProtT5 language model [15] (*ProtT5*) had overall similar results, demonstrating that the alphabetisation architecture and paradigm can in theory be applied to any current or future language model. Finally, as shown in Figure 2F, 20 is a reasonable choice for alphabet size, trading only minor performance increase for a more universally usable alphabet that can be straightforwardly plugged in to a number of protein sequence bioinformatics tools.

#### 2.2.2Fast and sensitive sequence search with TEA

To enable using TEA as a fast search tool with meaningful E-values, we developed sTEAm (Search with TEA against Many) which combines TEA and amino acid substitution scores during alignment, and uses exact *k*-mer matching and a frequency-aware TEA *k*-mer prefilter to boost search speed, and outputs calibrated E-values using a log-linear model as described in [23] (see Section 6.6 for details). As shown in Supplementary Section A.1, sTEAm provides better calibrated E-values than Foldseek. Overall, sTEAm delivers results with faster algorithms than typical amino acid searches at structure-level sensitivity (see Supplementary Table S2). We evaluated sTEAm to the MMseqs2 multidomain benchmark [1], which contains full length proteins, query sequences with scrambled regions, as well as reversed target sequences, thus also allowing a true test of the impact of sequence context on TEA sequence generation. Note that Foldseek cannot be used in this benchmark as structures for reversed and scrambled sequences are not well-defined. As shown in Supplementary Table S3, sTEAm greatly improves search sensitivity on this benchmark reaching sensitivity comparable to three iterations of MMseqs2 profile search and with improved speed. Furthermore, at this sensitivity, sTEAm returns much fewer false positives across fewer queries compared to profile search, where E-values based on BLOSUM62 become less reliable.

We developed a webserver (https://pickybinders.org/TEA/sTEAm) for multi-database searches, further described in Section 6.7.

### 2.3 Entropy as a measure of confidence and structure prediction

Associating predictions with reliable measures of confidence is very important, especially when using methods that are difficult to interpret, such as neural networks. A successful example is predicted LDDT (pLDDT), a metric provided by modern protein structure prediction methods like AlphaFold2/3 [3, 24] and ESMFold [14]. The pLDDT score estimates the local distance difference test (LDDT) [25] of a predicted model against a hypothetical ground truth, providing a per-residue confidence score. Previous research has shown that pLDDT can correlate both with the inability of the neural network to model the structure or with the presence of intrinsically disordered regions [26].

While we did not explicitly train a confidence predictor, the normalised Shannon entropy (Equation 2 in Section 6.2) associated with our model’s probability vectors can be interpreted as a measure of its certainty about a specific character. We therefore explored the normalized Shannon entropy of a probability vector as uncertainty at residue level and the average of the per-position entropies as uncertainty at sequence level. We observe that average entropy correlates inversely with search sensitivity in the SCOPe40 benchmark (Figure 3A), establishing it as a useful confidence measure for search results. Specifically, sequences with average entropy below 0.25 (*n*=2,729) achieved an average sensitivity of 0.81 *±* 0.29, whereas all sequences above this threshold (*n*=32) have zero sensitivity. In general, cases with low entropy (i.e. high confidence) but also low family sensitivity are typically difficult for both our method and Foldseek (Figure 3A bottom left, white), indicating that they represent challenging relationships for this benchmark.

**Fig. 3.**
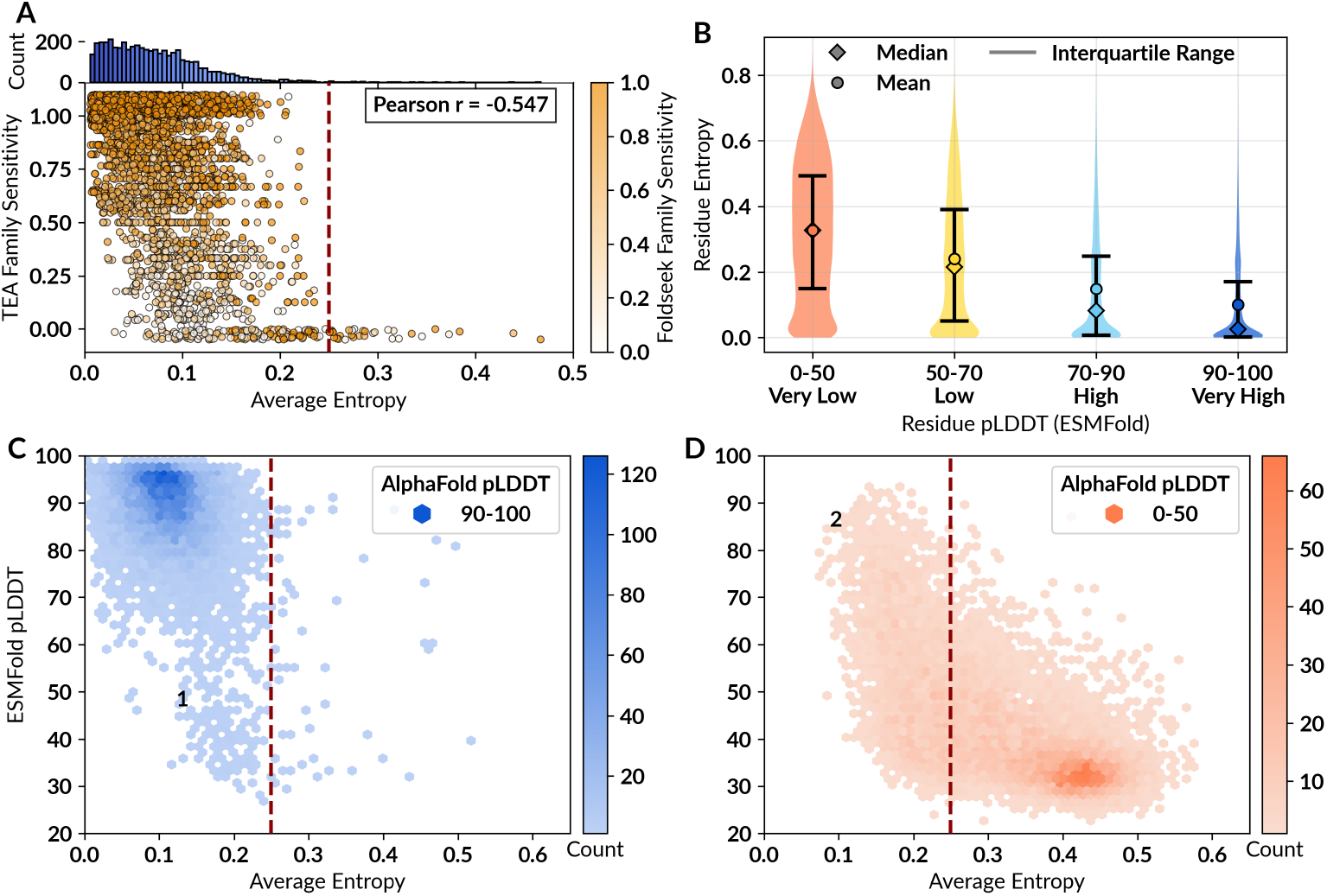
Entropy as a measure of confidence. **A)** Average TEA entropy vs. TEA family sensitivity up to the first FP from the SCOPe40 benchmark for all proteins from families of size 3 or more, coloured by the respective Foldseek family sensitivity. Points with family sensitivity of 0 and 1 have a small amount of jitter below 0 and above 1 respectively for visibility. **B)** Residue-level ESMFold pLDDT (binned) vs. residue-level TEA entropy for 40,000 residues from each pLDDT bin obtained from the 40k-pLDDT set (see Section 6.8). **C)** Average TEA entropy vs. average ESMFold pLDDT for 10,000 proteins where the corresponding average AlphaFold pLDDT is over 90. **D)** Same as C but for 10,000 proteins where the average AlphaFold pLDDT is less than 50. Points labelled 1 and 2 are described further in Supplementary Figure S4.

We also observed a significant correlation between TEA entropy and ESMFold pLDDT. Figure 3B shows the entropy distribution for residues from different pLDDT bins. We note that while the correlation trend is clear, many low pLDDT residues have low entropy, indicating that predicted disordered regions within otherwise high-confidence proteins are still consistently represented by our alphabet. Interestingly, we observed a strong correlation between average entropy and average pLDDT with a Spearman correlation of -0.823 to the maximum of AlphaFold and ESMFold pLDDT for the proteins in the 40k-pLDDT set (see Section 6.8). In Figure 3C-D, we show that entropy very often captures structural uncertainty, with high pLDDT proteins having low entropy (Figure 3C) and low pLDDT proteins having high entropy (Figure 3D). However, TEA is not limited to the accuracy of, for example, ESMFold which also uses pLM embeddings for structure prediction. We showcase two examples of such outliers with low TEA entropy in Supplementary Figure S4: the first case where the ESMFold pLDDT is low but AlphaFold pLDDT is high and second where the AlphaFold pLDDT is low and ESMFold pLDDT is high. In both cases, TEA search results against AFDB Clusters [27] and the PDB [12] demonstrate that the TEA sequence represents the higher pLDDT structure. This suggests that our alphabet can confidently and accurately model structural features even when structure prediction methods are unconfident.

### 2.4 Improving functional annotation with remote connections beyond AlphaFold2

Recently Foldseek was used to create the impactful AFDB Clusters dataset [27], which groups over 200 million proteins from AFDB into 15.3 million non-fragment clusters. However, only 2.3 million of these clusters have more than one entry - the remaining 13 million are singletons. We used TEA to connect these singletons to cluster representatives where possible. We used an average entropy threshold of *<*0.25 to select proteins where our alphabet would provide confident results (Figure 4A). This procedure filtered the dataset to 1.86 million representatives (81%) and only 5.24 million singletons (40%), indicating that many singletons are structurally and evolutionarily ambiguous both to Foldseek and pLMs, potentially pointing towards sequencing errors and protein fragments.

**Fig. 4.**
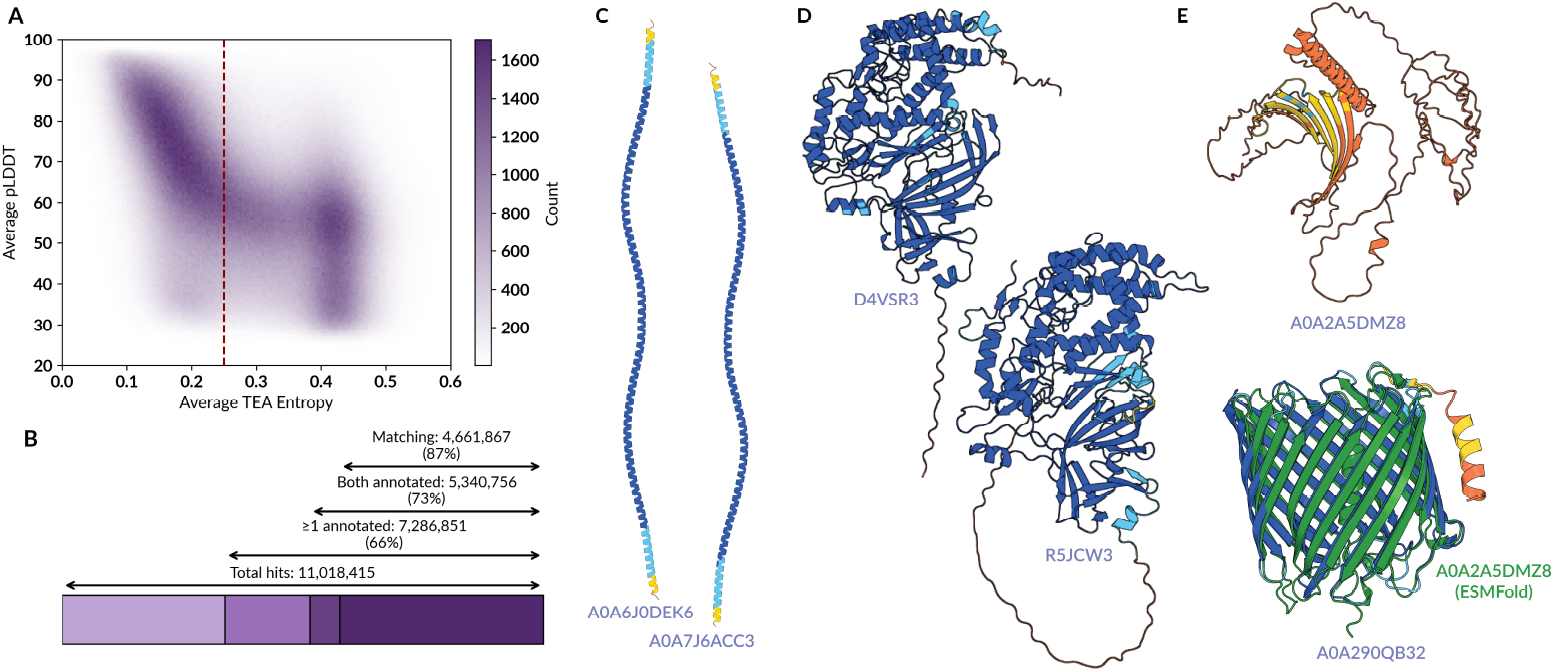
Remote homology in structural singletons. **A)** pLDDT vs.TEA entropy for all 15.3 million AFDB cluster representatives. The chosen entropy threshold is shown as a dashed red line. **B)** The total number of found hits at *>*90% coverage and E *<* 0.01 when searching singletons against all representatives (both with entropy *<*0.25) with sTEAm, the number/percentage of hits where either query and target have annotations in InterPro [28], where both have annotations, and where at least one annotation matches between the two. Percentages reported are always with respect to the previous count. **C-E)** Examples of singletons and their closest found cluster representative, missed by Foldseek. **C)** Proteins with mainly helix content **D)** Proteins with disordered regions. **E)** Proteins with low-confidence AlphaFold2 models. Note that for A0A2A5DMZ8, the ESMFold [14] model (shown in green) is a *β*-barrel with *<*3Å RMSD to the AFDB model of the closest hit, supporting the structural similarity suggested by TEA.

Searching with sTEAm with a 90% coverage threshold and E < 0.01 (as used in the original AFDB clustering) successfully found over 11 million hits for 545,495 singletons. Of these new connections, 48% had InterPro annotations for both query and target, and 87% of those annotations matched exactly (Figure 4B), demonstrating the high accuracy of our hits. Excitingly, over 1.95 million hits have either query or target annotated, but not the other, pinpointing cases where functional annotation transfer could be performed. Figure 4C-E shows examples of newly connected singletons and their closest cluster representatives, all of which have exact InterPro matches. These examples highlight three common use cases where our method is effective: (1) proteins with mainly helix content, where Foldseek often struggles to find meaningful structural matches due to the over-representation of 3Di-v (see Figure 1C), but the underlying sequences and embeddings still show detectable structural homology (panel C), (2) proteins with disordered regions which cause shorter structural alignments failing the 90% coverage threshold (panel D), and (3) incorrectly modelled proteins where a structural search would fail (panel E). In the latter case, a comparable structure could be produced with ESMFold [14].

In summary, TEA generates highly relevant and complementary comparisons to structure-based methods while requiring minimal computational resources (<800 MB storage for >10 mil. sequences, and just 2.5h for 5.2M × 7.1M comparisons with 64 cores and 64 GB RAM). This proof of concept demonstrates that TEA similarities could be used with appropriate coverage and E-value thresholds, perhaps even in combination with sequence and structure similarities, to recreate clustering efforts such as AFDB Clusters [27] and UniProt3D [29] from a novel perspective, bringing new functional connections and discoveries to both domain-level and protein-level analyses.

## 3 Discussion

We developed a new alphabet derived from protein language models that allows the identification of remote homologs with performance comparable to structure-based methods, along with a new search service using this alphabet. TEA offers several key advantages over existing protein sequence representations. As with standard amino acids, highly optimized and efficient sequence tools such as MMseqs2 can be applied, but, unlike amino acids, it facilitates the identification of remote structural homologs and enables motif discovery. TEA shows better inherent structural representation capability than Foldseek’s 3Di despite requiring no prior structural information, and ultimately complements structural representations by being more effective at comparing predicted disordered and repetitive regions. We use the inherent capabilities of the alphabet to power sTEAm (Search with TEA Against Many), a search tool that enables exceptionally fast sequence search while maintaining the ability to detect highly remote homology. This capability can be used to generate query-centric MSAs for structure prediction, potentially improving the modelling of proteins that currently suffer from shallow MSAs with traditional search methods. Furthermore, pairing this tool with profile-generation methods like HMMER [30] might improve the sensitivity and quality of sequence profiles for broader evolutionary analysis, or help with structural phylogenetics [31]. Beyond search applications, the TEA representation and its associated entropy, which correlates with pLDDT, offer compelling opportunities for *de novo* protein design. Because these metrics provide direct insight into fold plausibility without relying on computationally expensive structure prediction methods, they enable extremely fast Monte Carlo sampling, as recently explored in [32]. Furthermore, deeper analysis of TEA entropy could highlight regions of protein sequence space that are difficult for protein language models (pLMs) to represent, ultimately guiding the improvement of future pLMs and embedding representation.

Beyond TEA’s immediate utility, the method provides a generalisable framework for training specialised alphabets from pLM embeddings using a contrastive objective. This architecture is readily adaptable, allowing researchers to fine-tune alphabets for specific goals. For instance, an alphabet could be trained for function prediction with characters grouping residues by their functional roles (e.g., active site, binding pocket) rather than purely structural characteristics. Another possibility is to develop an alphabet for interface description, enabling the clear characterisation of protein-protein interaction interfaces, which are often subtle and complex in continuous space. Alternatively, researchers could apply the same discretisation and contrastive learning principles to RNA sequences and their high dimensional descriptors [33], enabling highly efficient searches that might improve modelling efforts [34].

Ultimately, TEA brings the powerful representation capabilities of deep learning to well-established sequence bioinformatics algorithms, such as homology search, profiles, phylogenetic trees, motif finding, multiple sequence alignments, and more, all while maintaining the speed and low resource consumption of amino acid sequences.

## 4 Code and data availability

The TEA model code, sequence conversion scripts, training scripts, and documentation are provided at github.com/PickyBinders/TEA. TEA is also available on Hugging Face at huggingface.co/PickyBinders/TEA. The sTEAm search tool is available at https://github.com/PickyBinders/sTEAm. An online search server is available at https://pickybinders.org/TEA and also provides TEA FASTA files of the following converted databases: AFDB Clusters (v3), UniRef50 (version 2026_01). Benchmarking data are available at https://doi.org/10.5281/zenodo.17725635.

## 5 Acknowledgements

We thank the members of the Schwede group for insightful discussions and technical support, and sciCORE at the University of Basel (https://scicore.unibas.ch/) for providing computational resources and storage space. We gratefully acknowledge financial support for parts of this work by the SIB Swiss Institute of Bioinformatics (https://www.sib.swiss/), the Biozentrum of the University of Basel (https://www.biozentrum.unibas.ch/), and the Swiss National Science Foundation (SNSF; Ambizione grant 223634 for JD, WEAVE grant 220141 for LP).

## 6 Methods

### 6.1 Training data

Domains from SCOPe40 (v2.08) [35] were structurally aligned using TM-align and all alignments from within the same SCOPe40 superfamily with TM-score above 0.6 were retained. To generate triplets of residues, all aligning residue pairs with *Cα* RMSD below 5 Å were considered as anchor and positive, with corresponding negatives for each anchor being randomly selected from a 5-residue window around the aligning position. The embeddings were obtained from the 4-bit quantized ESM2-650M model (`facebook/esm2_t33_650M_UR50D on HuggingFace`) [36].

#### 6.1.1 Cross-validation

To assess generalisation potential, we also trained versions of TEA by four-fold cross-validation on SCOPe40, as described previously for Foldseek [6]. The SCOPe40 dataset is divided into four parts, such that all domains of each fold ended up in the same part of the four parts. The training data triplets were thus split into these folds and alphabets were trained on three parts and tested on the remaining part, selecting each of the four parts in turn as a test set. As good generalisation was seen in the cross-validation experiment, the final TEA model is trained on the entire training set.

### 6.2 Model training

The embeddings from `facebook/esm2_t33_650M_UR50D` were discretized into an alphabet of 20 characters using contrastive learning on residue triplets, on an architecture consisting of a dense linear layer (1280 × 1280), a layer normalisation layer, a GELU activation function, and the final linear layer (1280 × 20) resulting in a model with 1.7 million trainable parameters. The model was trained for 10 epochs with the AdamW optimiser [37] (weight decay 0.3), a cosine annealing learning rate scheduler (from initial learning rate 0.005 to 0.0001), and a dropout of 0.1 added after the activation function.

The main, contrastive objective was to minimize the similarity between the anchor and the negative sample while maximizing the similarity between the anchor and the positive sample in each triplet. This approach inherently encourages the alphabet to represent increased similarity for aligning residues and decreased similarity for non-aligning ones. Given **a, p** and **n** the probability vectors generated by our model for respectively the anchor, positive, and negative, the contrastive loss was implemented as follows:

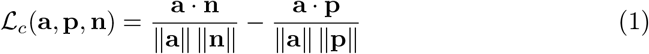

We also implemented two auxiliary losses: a Shannon entropy loss and a uniform loss. Lower entropy indicates a prediction strongly committed to a specific character, whereas higher entropy suggests the model is less certain about its character choice. We minimized the Shannon entropy associated to the probability vectors of the triplets with *N* being the logits dimension (alphabet size):

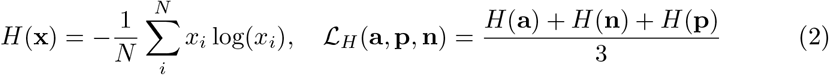

To encourage uniform character usage, the uniform loss was achieved by applying a Kullback-Leibler (KL) divergence loss against a uniform distribution, **u**. For each batch we averaged the probabilities relative to the predicted logits into a vector **b** and we computed a KL divergence loss as follows:

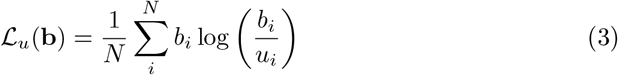

The final loss of the model was weighted as follows:

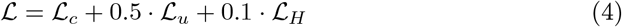

#### 6.2.1 Ablations

To assess the impact of model and training choices, we trained a number of models with different settings ablated. These include *“No uniform loss”* and *“No entropy loss”* models where the ℒ_*u*_ and ℒ_*H*_ terms respectively in Equation 4 are removed, a *“Far window”* model where the negatives for each triplet were selected randomly from a window of 5-10 residues away from the aligning position insTEAd of 1-5, a *“Full precision”* model where 4-bit quantisation is not used for embedding generation, and *“ESM-3B”* and *“ProtT5”* models where the pLMs used for embedding generation are changed to the `facebook/esm2_t36_3B_UR50D` model and the `Rostlab/prot_t5_xl_uniref50` model respectively, with corresponding changes in input embedding dimension. All of these ablations are trained on the 4-fold SCOPe fold split. Further, models with different alphabet sizes (4, 8, 12, 16, 24, 28, 32, and 40) were trained on the full training set.

### 6.3 TEA substitution matrix

We created a BLOSUM-like substitution matrix for TEA sequences from pairs of structurally aligned residues used for training. First, we determined the TEA states of all residues. Next, the substitution frequencies among TEA states were calculated by counting how often two TEA states were structurally aligned. (Note that the substitution frequencies from state A to state B and the opposite direction are equal.) Finally, the score for substituting state *x* through state *y* is 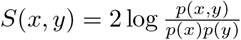.

### 6.4 Benchmarks

The SCOPe40 benchmark is defined and evaluated as described in [6], except applied to the latest version of SCOPe40 (v2.08) consisting of 15,128 single domains with a median length of 149 residues. The SCOPe40c dataset is a curated subset of SCOPe v1.75 with 9,705 domains, described in [23]. The family and superfamily benchmark plots measure the sensitivity of detected TPs (same family, and same superfamily but not same family respectively) up to the first FP (where FP is defined as hit from a different fold). For 4-fold cross-validation results (labelled as CV in the figures), the sensitivity curves and AUCs are calculated only for comparisons between domains in the held-out folds. For all methods, each query was allowed to have up to 2000 hits (controlled by the`–max-seqs` parameter in MMseqs2 and Foldseek).

The reference-free multidomain benchmark is adapted from [6] - we aligned all 688,852 community representatives from UniProt3D [29] using BLAST (2.5.0+) with an E-value threshold *<* 10^*−*3^ and then clustered using SPICi [38], resulting in 56,574 clusters. For each cluster, we picked the longest protein as representative. We randomly selected 100 representatives as queries and searched the set of remaining proteins. Foldseek and MMseqs2 searches were run as described in [6], while TEA search was run using the *exact* mode described in Section 6.5. Evaluation was performed using the scripts obtained from github.com/steineggerlab/foldseek-analysis.

The MMseqs2 multidomain benchmark is defined in [1], as containing 6,370 query and 30.4M target sequences all derived from full-length UniProt sequences, and each annotated with structural domain annotations from SCOP. 27M of the target sequences are reversed, and unmatched parts of query sequences are scrambled in a way that conserved the local amino acid composition. True-positive matches are defined as those that have annotated SCOP domains from the same SCOP family; false positives match a reversed sequence or a sequence with a SCOP domain from a different fold. Other cases are ignored.

### 6.5 Alignment methods

TEA is designed to be compatible with any classical alignment tool. In this work, we used MMseqs2 (version 18.8cc5c) [1] to perform alignments across three distinct modalities detailed below. When indicated as *sensitive*, we run the standard MMseqs2 easy-search command as follows:

~~~
mmseqs easy-search TEA_query.fasta TEA_target.fasta results.m8 tmp/
--comp-bias-corr 0 --mask 0 --gap-open 18 --gap-extend 3
--sub-mat matcha.out --seed-sub-mat matcha.out
~~~

For the *exact* modality, we include the flag `–exact-kmer-matching` 1 in the easy-search command. This modification significantly increases the speed of the command by restricting the pre-filtering step to only allow exact *k*-mer matches. For 3Di and amino acids, gap open and gap extend penalties of 12 and 1 were used, as these gave the best results on a grid-search for the SCOPe benchmark.

The sequences from models with different alphabet sizes (Figure 2D) and alphabet combinations (Figure 2A, Supplementary Figure S2) insTEAd use all vs. all alignments with no pre-filtering step, followed by taking the top 2000 alignments per query (labelled *exhaustive*). These alignments were calculated using Needleman-Wunsch-style dynamic programming with gap penalties on score matrices, constructed such that *M*_*ij*_ represents the substitution score of residue positions *i* and *j*. We used BLOSUM62 for amino acids, the 3Di substitution matrix for 3Di, the TEA substitution matrix for TEA, and corresponding substitution matrices built as described in Section 6.3 for the different alphabet sizes. Combinations of alphabets use a matrix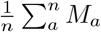where *M*_*a*_ is the score matrix constructed for alphabet *a* and *n* the number of alphabets combined. All alignments used gap open and extend penalties of 12 and 1.

### 6.6 Search with TEA against Many (sTEAm)

sTEAm is built on MMseqs2 [1] and adapts Foldseek’s dual-alphabet architecture [1, 6] by replacing the structure-derived 3Di alphabet with TEA, scoring Smith-Waterman alignments under both the TEA-specific substitution matrix (Section 6.3) and BLO-SUM62, with the amino-acid contribution upweighted by 1.4. We noted a TEA-specific prefilter pathology, namely a small number of recurrent structural motifs concentrates the index into very few *k*-mers that dominate prefilter compute without contributing discriminative signal (on SCOPe40c the top 0.1% of *k*-mers hold *∼*20% of all index entries and account for *∼*88% of prefilter lookups). Thus, sTEAm’s prefilter skips query lookups for any *k*-mer whose target bucket falls within the top 20% of the index by cumulative entry coverage. The sTEAm source code and package is available at https://github.com/PickyBinders/sTEAm.

#### 6.6.1 E-values

We developed an empirical E-value model for the combined TEA and amino acid scoring system in sTEAm. Standard Karlin-Altschul statistics assume a Gumbel-distributed score distribution, which does not hold for structural alphabet alignments due to convergent evolution of secondary and tertiary structure motifs producing a heavy-tailed false positive distribution [23]. InsTEAd, we adopted the proposed log-linear model where the cumulative false positive distribution *C* (*s*|*FP*) is approximated as *log*_10_(*C* (*s*|*FP*)) = *ms* + *c*, giving E-values of the form 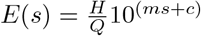, where *s* is the raw alignment score, 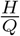is the average number of reported hits per query (computed at runtime from the prefilter output), and *m* and *c* are parameters fitted by log-linear regression of the empirical cumulative false positive distribution on the SCOPe40c dataset, a curated subset of SCOPe v1.75 with 9,705 domains [23]. False positives were defined as pairs from different SCOPe folds; pairs within the same fold but different superfamilies were excluded. The log-linear parameters were fitted on the FPEPQ (false positive errors per query) range of 1-10, yielding *m* = -0.018254 and *c* = -0.445 (*R*^2^ = 0.99). Because the log-linear model under-estimates the empirical false-positive rate at user-relevant thresholds we calibrate by modifying all reported E-values by a constant + log_10_ 3 *≈* 0.477 shift on the intercept, giving the final value for *c* = 0.032. sTEAm results are sorted by E-value (ascending), which corresponds to descending alignment score.

### 6.7 Web server

A website was developed to facilitate access to TEA-converted datasets, currently offering downloads in FASTA format for UniRef50 (version 2026_01) [39] and AlphaFold Clusters (v3) [27], with additional sources planned for future inclusion. Users may also submit amino acid sequences directly for conversion into TEA sequences (on CPU), which are queried against the available datasets using a simplified implementation of sTEAm. The search implementation follows the same general algorithmic frame-work as sTEAm (see Supplementary Table S1), but uses shorter prefilter *k*-mers of length 5 insTEAd of 6, enabling the full index to remain memory-resident for interactive queries. Results are queried via FastAPI [40], with results persisted in MongoDB [41]. The interface is built with Next.js [42], and where a search hit has a known 3D structure, it is fetched from the MongoDB and visualised interactively using the NGL structure viewer [43]. The underlying search implementation is available at https://git.scicore.unibas.ch/schwede/kmatch.

### 6.8 40k-pLDDT set

Sequences and 3Di characters for the character comparison analysis were obtained from the SCOPe40 domains, and from 40,000 proteins with lengths between 100 and 600 selected randomly from the Foldseek `alphafold_uniprot50` database such that 10,000 proteins each had an average AFDB pLDDT within the bins 0-50 (“Very low”), 50-70 (“Low”), 70-90 (“High”) and 90-100 (“Very high”). ESMFold [14] structures were created for all proteins in this set.

### 6.9 AFDB clusters

We obtained the UniProt accessions corresponding to AFDB cluster v3 representatives from the `5-allmembers-repId-entryId-cluFlag-taxId.tsv` file in afdb-cluster.steineggerlab.workers.dev and restricted representatives to those with cluFlag set to 2 (singletons) and 4 (non-singleton representatives). These sequences were converted to TEA sequences and filtered with an average TEA entropy threshold of 0.25. The sTEAm search was run for the singletons passing the threshold against all sequences passing the threshold with a coverage threshold of 90% across both query and target, and an E-value threshold of 0.01.

## Appendix A Supplementary information

**Supplementary Figure S1.**
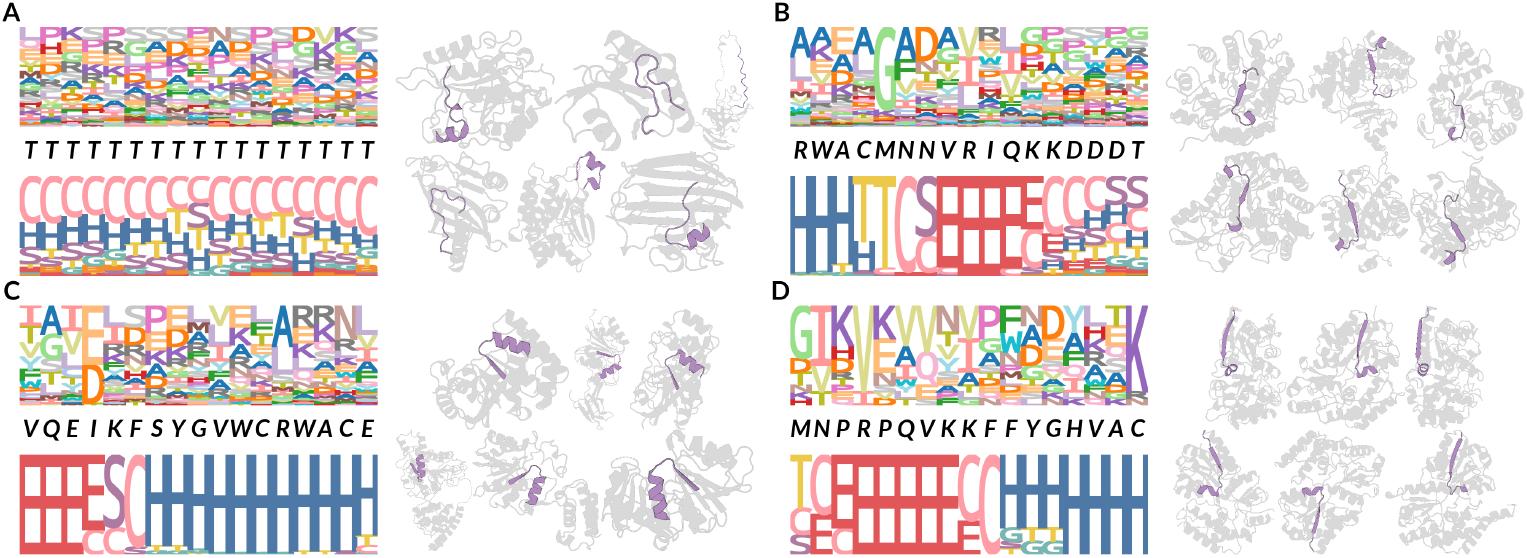
Conserved TEA motifs. Examples of TEA motifs of length 17 found in 76 SCOPe40 IDs across 56 different folds, **B)** in 261 IDs across 26 superfamilies only in fold c.1 **C)** in 79 IDs across 40 families only in superfamily c.66.1, and **D)** 17 IDs only in family c.94.1.0 are shown. For each motif, the left panel shows the amino acid frequency logo, the TEA characters of the motif in italics, and the DSSP frequency logos for all occurrences in SCOPe40, and the right panel shows examples of structures (gray) containing the motif (highlighted in purple). DSSP labels are as follows, H: *α*-helix, G: 3-10 helix, E: *β*-sheet, C: random coil, T: H-bonded turn, S: bend.

**Supplementary Figure S2.**
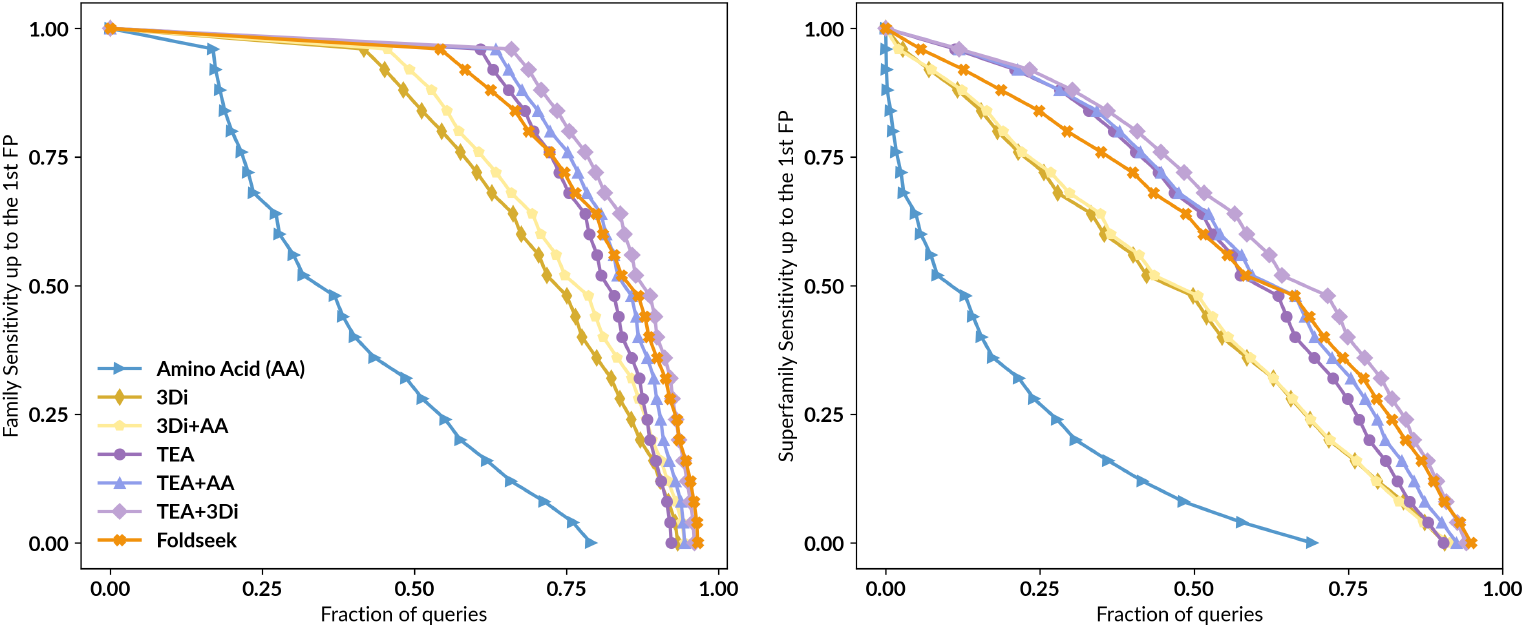
Exhaustive alignments of alphabet combinations. **A)** Cumulative distributions of sensitivity for homology detection on the SCOPe40 database of single-domain structures. TPs are matches within the same family; FPs are matches between different folds. Sensitivity is the area under the ROC curve up to the first FP. **B)** Same as A with TPs as matches within the same superfamily.

### A.1 E-value calibration

We evaluated the effect of E-value calibration on homology detection sensitivity using an all-against-all search on SCOPe40c (a curated subset of SCOPe40 [23]) with `–max-seqs` set to 2000. As seen in Table S1, without any E-value filtering, sTEAm achieved sensitivities comparable to Foldseek and to TEA-only search using MMseqs2. When applying a threshold of *E <* 10, sTEAm sensitivity remained high at an empirical false-positive rate of 23.6, confirming the calibration accuracy of the log-linear model. Foldseek’s native E-values, by contrast, substantially overestimate significance: at its reported *E <* 10 threshold the empirical false positive errors per query (FPEPQ) is 441.4, consistent with previous observations that structural-alphabet E-values based on Gumbel distributions fail to account for the heavy-tailed false-positive score distribution arising from convergent structural evolution [23]. To enable a fair comparison, we determined the Foldseek E-value threshold at which the empirical FPEPQ matched sTEAm’s rate of 23.6 and filtered Foldseek’s results accordingly. At matched false positive rates, sTEAm’s calibrated E-values provide comparable sensitivity to Foldseek while offering meaningful statistical significance estimates.

**Supplementary Table S1.**
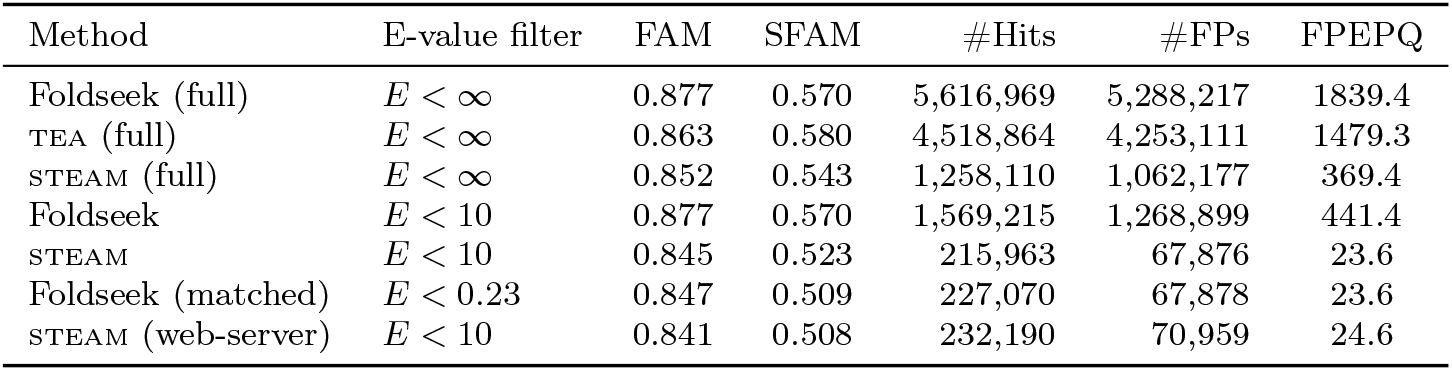
E-value calibration. Homology detection sensitivity on SCOPe40c (*N* = 2,875 evaluable queries). FAM and SFAM report mean family and superfamily sensitivity before the first false positive (different fold). FPEPQ is the empirical false positive errors per query.

| Method | E-value filter | FAM | SFAM | #Hits | #FPs | FPEPQ |
| --- | --- | --- | --- | --- | --- | --- |
| Foldseek (full) | $E < \infty$ | 0.877 | 0.570 | 5,616,969 | 5,288,217 | 1839.4 |
| TEA (full) | $E < \infty$ | 0.863 | 0.580 | 4,518,864 | 4,253,111 | 1479.3 |
| STEAM (full) | $E < \infty$ | 0.852 | 0.543 | 1,258,110 | 1,062,177 | 369.4 |
| Foldseek | $E < 10$ | 0.877 | 0.570 | 1,569,215 | 1,268,899 | 441.4 |
| STEAM | $E < 10$ | 0.845 | 0.523 | 215,963 | 67,876 | 23.6 |
| Foldseek (matched) | $E < 0.23$ | 0.847 | 0.509 | 227,070 | 67,878 | 23.6 |
| STEAM (web-server) | $E < 10$ | 0.841 | 0.508 | 232,190 | 70,959 | 24.6 |

**Supplementary Table S2.** Runtime and performance for SCOPe40 (15,128 proteins). The max-seqs column describes the number of hits allowed to pass the pre-filtering step. EBA does not have a pre-filtering step and thus all pairs of alignments are computed and then sorted to take the top 1000 per query. All methods were run with 64G RAM and 128 cores. Note that Foldseek times reported do not include database creation time.

| Method | max-seqs | Time (s) | Family AUC | Superfamily AUC | Fold AUC |
| --- | --- | --- | --- | --- | --- |
| MMseqs2 | 300 | $3.03 \pm 0.08$ | 0.214 | 0.081 | 0.000 |
| | 1000 | $3.04 \pm 0.02$ | 0.214 | 0.081 | 0.000 |
| STEAM | 300 | $3.39 \pm 0.01$ | 0.795 | 0.553 | 0.053 |
| | 1000 | $5.24 \pm 0.03$ | 0.801 | 0.561 | 0.058 |
| Foldseek | 300 | $25.18 \pm 0.03$ | 0.764 | 0.512 | 0.075 |
| | 1000 | $39.45 \pm 0.19$ | 0.805 | 0.560 | 0.095 |
| EBA | - | 7,560 | 0.84 | 0.6 | 0.079 |

**Supplementary Figure S3.**
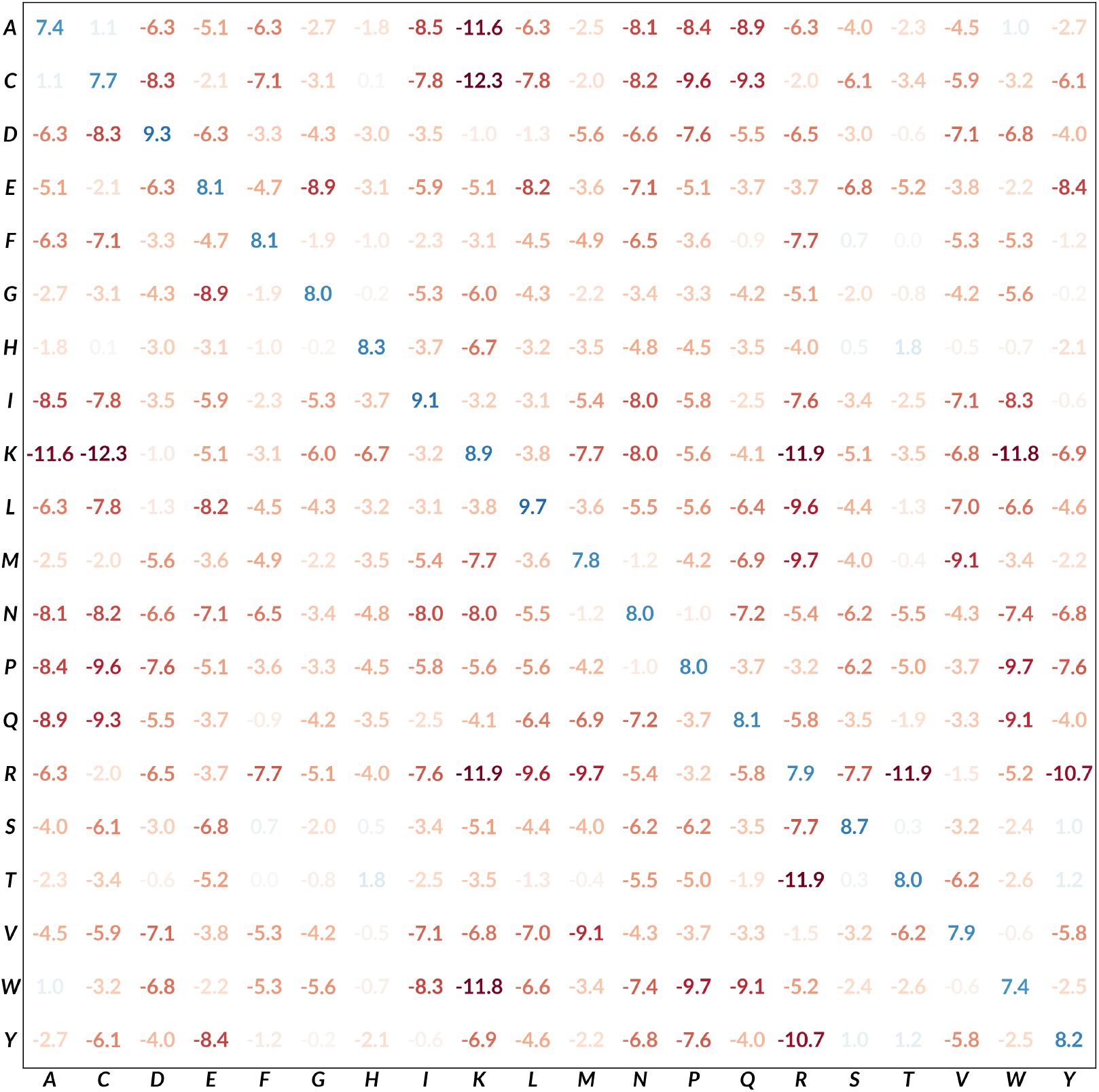
TEA substitution matrix. See Section 6.3 for how the matrix is computed. The numbers are colored on a **red-blue** color scale with **red** being the lowest and **blue** the highest.

**Supplementary Figure S4.**
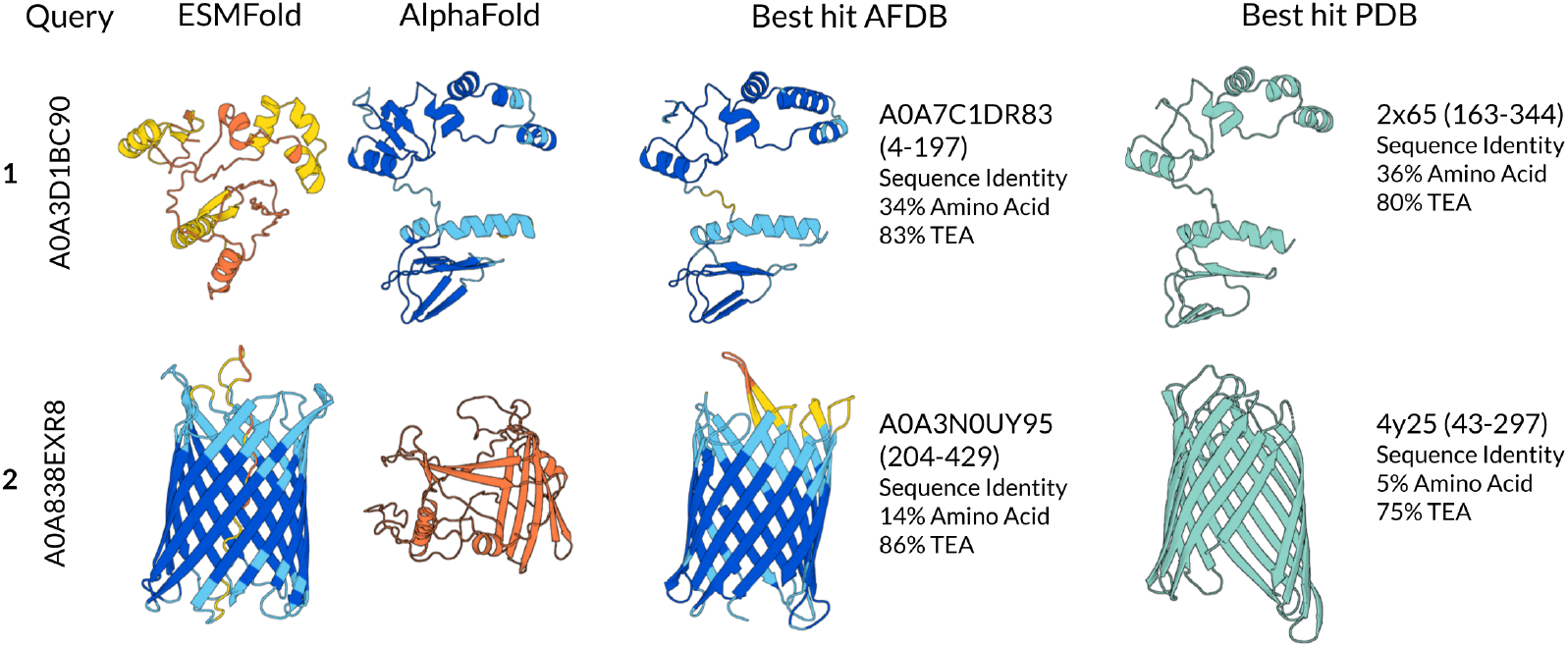
TEA confidently represents the correct structure even in AlphaFold and ESMFold failure modes. Two examples of searches for proteins highlighted in Figure 3C-D, displayed as the ESMFold and AlphaFold structures of the protein respectively, and then the closest TEA hit for the protein in AFDB Clusters and in the PDB respectively, both cropped to the alignment region with residue range displayed. The amino acid and TEA sequence identities of the TEA alignments are also displayed.

**Supplementary Table S3.** MMseqs2 multidomain benchmark. Analysis of search sensitivity (TP = same family, FP = different fold), high-scoring false positives, and runtime for 6,370 searches with UniProt sequences through a database of 30 million UniProt-derived sequences.

| Method | AUC up to first false positive | Fraction of queries with false positive at $E < 10^{-3}$ | Reported false positives at $E < 10^{-3}$ | Runtime (s) | # of alignments returned |
| --- | --- | --- | --- | --- | --- |
| MMseqs2 | 0.329 | 0.004 | 520 | 266s | $5 \times 10^6$ |
| MMseqs2 3 IT | 0.637 | 0.134 | 269,179 | 577s | $9 \times 10^6$ |
| STEAM | 0.623 | 0.016 | 67,875 | 219s | $25 \times 10^6$ |

## Footnotes

1 TEA characters are shown in italics and 3Di in lowercase; the amino acid letters are used for depiction purposes but no direct amino acid correspondence is expected for either alphabet.

